# Abundance and nuclear antigen reactivity of intestinal and fecal Immunoglobulin A in lupus-prone mice at younger ages correlate with the onset of eventual systemic autoimmunity

**DOI:** 10.1101/2020.06.25.172155

**Authors:** Wei Sun, Radhika Gudi, Benjamin M. Johnson, Chenthamarakshan Vasu

## Abstract

Our recent studies, using (SWRxNZB)F1 (SNF1) mice, showed a potential contribution of the gut microbiota and pro-inflammatory immune responses of the gut mucosa to systemic autoimmunity in lupus. Here, using this mouse model, we determined the abundance and the nAg reactivity of IgA antibody produced in the intestine under lupus susceptibility. Intestinal lymphoid tissues from SNF1 mice, females particularly, showed significantly higher frequencies of nAg (dsDNA and nucleohistone) reactive IgA producing B cells compared to B6 females. Most importantly, younger age fecal IgA -abundance and - nAg reactivity of lupus-prone mice showed a positive correlation with eventual systemic autoimmunity and proteinuria onset. Depletion of gut microbiota in SNF1 mice resulted in the diminished production of IgA in the intestine and the nAg reactivity of these antibodies. Overall, these observations show that fecal IgA features, nuclear antigen reactivity particularly, at preclinical stages/in at-risk subjects could be predictive of autoimmune progression.

## 1. Introduction

Systemic lupus erythematosus (SLE) is an autoimmune disease which arises when abnormally functioning B lymphocytes, in at risk subjects, produce auto-(self-reactive) antibodies to nuclear antigens such as DNA and proteins. High levels of circulating autoantibodies and immune complex deposition in the kidney, leading to tissue damage and glomerulonephritis are the hallmarks of SLE ^1^. Importantly, women are more predisposed to SLE than men, and the disease prevalence ratio of women is about 9:1 over men ^2^. Autoantibody production and gender bias in SLE is caused by a combination of genetic and environmental factors ^1-4^. Disproportionate functioning of genes as well as sex hormones, estrogen in particular, contribute to the onset and development of disease activities in SLE ^2,5-8^. Recent studies that used human samples and rodent models have shown that gut microbiota composition influences the rate of disease progression and the overall disease outcome ^9-15^. We have demonstrated that minor dietary deviations alter the composition of gut microbiota and SLE in a mouse model ^13^. We have also found that gut microbiota influences the autoimmune progression differently in lupus-prone male and female mice, leading to a gender bias in disease incidence^16^.

Our recent studies that used lupus-prone (SWRxNZB)F1 (SNF1) mice showed a potential contribution of pro-inflammatory immune response initiated in the gut mucosa, and gut microbiota in triggering the disease associated gender bias observed in SLE ^16,17^. We also showed that pro-inflammatory responses including higher cytokine expression, recruitment of large number of immune cells, and presence of higher number of antibody positive plasma cells in the gut mucosa of lupus-prone females, compared to males, can be detected as early as at juvenile age. These pro-inflammatory immune features of female mouse gut mucosa progressively increase at later ages, prior to systemic autoimmunity and kidney pathology. These observations and reports by others showing the involvement of microbiota in systemic autoimmune progression in lupus ^10-12,18,19^ suggest that autoantibody production and systemic autoimmunity in lupus-prone subjects are initiated in the gut mucosa, microbiota dependently and there is a need for additional studies to assess antibody production in the intestine.

IgA is the most abundant Ig isotype released in to the gut lumen and it plays an important role in the protection against microbial infection as well as in maintaining a healthy gut microbiota ^20-22^. Intriguingly, a recent report showed, in addition to differences in the gut microbiota composition, relatively higher levels of total IgA in stool samples of SLE patients compared to that of healthy controls ^9^. On the other hand, serum IgA levels, but not IgG or IgM levels, were diminished in lupus-prone mice that received oral treatment with Lactobacillus, which suppresses lupus nephrites ^23^. Importantly, anti-DNA antibodies of IgA class are found in the serum of patients with SLE ^24- 29^, suggesting that they may be of gut primed B cell origin. These reports along with our studies ^16,17^ showing pro-inflammatory immune phenotype and higher plasma cell frequency by lupus-prone female mouse intestine suggests the degree of IgA secretion in the gut lumen could show gender bias and may be indicative of lupus susceptibility and autoimmune progression. Nevertheless, the relationship between fecal IgA levels and gender bias in lupus is unknown. Further, the reactivity of fecal IgA in a lupus-prone background with nuclear antigens and the potential association with disease onset has never been studied.

In the present study, we investigated the degree of IgA production in the intestine, and the abundance and nAg reactivity of fecal IgA in lupus-prone SNF1 mice. We have then assessed the relationship between these features and autoimmune progression in male and female mice, and if the gut microbiota has an influence on fecal IgA abundance and nAg reactivity. Our studies, for the first time, show not only that higher amounts of IgA are produced in the gut mucosa-associated immune tissues of lupus-prone mice, but also these antibodies have significant nAg reactivity, even at juvenile age. Correspondingly, the abundance of IgA is profoundly higher in the feces of lupus-prone mice, females particularly, and these antibodies show significant nAg (dsDNA and nucleohistone) reactivity. These fecal IgA features of pre-clinical stages correlate significantly with the eventual systemic autoantibody levels and the proteinuria onset in lupus-prone mice, suggesting that fecal IgA features could be valuable for predicting systemic autoimmune progression, early on, prior to sero-positivity and lupus nephritis.

## 2. Materials and methods

### 2.1 Mice

SWR/J (SWR), NZB/BlNJ (NZB), C57BL/6J (B6), and Balb/cJ mice were purchased from the Jackson Laboratory (Bar Harbor, Maine) and housed under specific pathogen free (SPF) conditions at the animal facilities of Medical University of South Carolina (MUSC). (SWRxNZB)F1 (SNF1) hybrids were generated at the SPF facility of MUSC by crossing SWR females with NZB males. Fresh fecal pellets from collected separately from individual mice at different time-points (SNF and B6 mice; bred in our facility). Fecal pellets from Balb/cJ mice were collected immediately upon receipt from the vendor. In some experiments, SNF1 mice were given a broad spectrum antibiotic cocktail as described in our recent report^30^ to deplete the gut microbiota. Briefly, female SNF1 mice were given a broad-spectrum antibiotic cocktail [ampicillin (1 g/l), vancomycin (0.5 g/l), neomycin (1 g/l), and metronidazole (1 g/l)] -containing drinking water, starting at week 4 of age. Depletion of gut microbiota was confirmed by the culture of fecal pellet suspension on blood agar and brain heart infusion agar plates, under aerobic and anaerobic conditions, as described before^30^. Urine and tail vein blood samples were collected at different time-points to detect proteinuria and autoantibodies. All animal experiments were performed according to ethical principles and guidelines were approved by the institutional animal care committee (IACUC) of MUSC. Animal experimental protocol of this study was approved by the IACUC of MUSC.

### 2.2 Proteinuria

Urine samples were tested weekly for proteinuria. Protein level in the urine was determined by Bradford assay (BioRad) against bovine serum albumin standards as described before^13,16,17^. Proteinuria was scored as follows; 0: 0-1mg/ml, 1: 1-2mg/ml, 2: 2-5 mg/ml, 3: 5-10mg/ml and 4: ≥ 10mg/ml. Mice that showed proteinuria >5 mg/ml were considered to have severe nephritis.

### 2.3 Immune cell culture

Male and female SNF1 mice, and female B6 mice of 4 and 16 weeks of age were euthanized, single cell suspensions of spleen and Peyer’s patches (PP) or enriched immune cells from ileum portion of the small intestine (Si) and large intestine (Li) were prepared as described in our recent reports ^13,16,17^. These cells were subjected to flow cytometric analysis to detect IgA+ B cells as described before ^17^, and/or cultured in the presence of bacterial LPS (2 μg/ml) and anti-CD40 antibody (5 μg/ml) for 48 h. Optimally diluted, spent media were tested for relative levels of total IgA and nucleohistone- and dsDNA-reactive IgA by ELISA.

### 2.4. ELISA

Antibodies against nAgs (nucleohistone and dsDNA) in mouse sera were evaluated by ELISA as described in our recent reports with minor modifications ^13,17^. Briefly, 0.5μg/well of nucleohistone (Sigma-Aldrich) or dsDNA from calf thymus (Sigma-Aldrich) was coated as antigen, overnight, onto ELISA plate wells. Serial dilutions of the sera were made, total IgG against these antigens were detected using HRP-conjugated respective anti-mouse antibody (Sigma-Aldrich, eBioscience and Invitrogen) and the reaction was detected using TMB substrate (BD biosciences). For determining antibody levels in fecal samples, extracts of fresh fecal pellets from individual mice were collected separately and used. Weighed fecal pellets were suspended in proportionate volume of PBS (5% suspension or 50 mg feces/ml; w/v) by breaking the pellet using a pipette tip and high speed vortex, and continuous shaking at 800 rpm overnight at 4°C. Suspensions were centrifuged at 14,800 rpm for 15 min and top 2/3 of the supernatants were collected. Supernatants were diluted 1:100 for determining total IgA concentration. Total IgA levels were determined by employing in-house quantitative sandwich ELISA. Briefly, purified anti-mouse IgA monoclonal antibody (0.1 μg/well in 50 μl) coated wells were incubated with diluted samples for 2 h, incubated with biotin linked polyclonal anti-mouse IgA antibody for 1 h, and finally with avidin-HRP for 30 min, before developing the reaction using TMB substrate. Purified mouse IgA (Southern biotech) was used in each plate for generating the standard curve. In initial assays, to test anti-dsDNA and anti-nucleohistone reactivity of IgA in different sample types, optimum dilution (1:10) of aforementioned extracted samples were incubated in dsDNA or nucleohistone coated plates and further incubated with anti-mouse IgA-HRP before developing the assay. In ELISA assays where dsDNA or nucleohistone antibody titers were determined, serial dilutions of fecal extracts (starting at 1:10 dilution) were incubated in dsDNA or nucleohistone coated plates, incubated with biotin linked anti-mouse IgA antibody, and followed by avidin-HRP. Known positive and negative control samples identified from the initial screening were used in all plates to validate the results for determining a reliable titer. Highest dilution of the sample that produced an OD value of ≥0.05 above background value was considered as the nAg reactive titer. IgA concentrations and nAg reactive titers per gram of feces were then calculated for the data presented here.

### 2.5 Statistical analysis

GraphPad Prism or Microsoft excel was used to calculate the statistical significance. Two-tailed *t*-test or Mann-Whitney test was employed to calculate *p*-values when two means were compared. Proteinuria scores were analyzed using the Chi-square test. Pearson’s correlation coefficient/bivariate correlation approach was employed for measuring linear correlation between two variables. For correlating concentration and titer of two specific factors, Spearman correlation approach was employed. A *P* value ≤0.05 was considered statistically significant.

## 3. Results

### 3.1 Higher amounts of IgA are produced in the gut mucosa of lupus-prone SNF1 mice

SNF1 mice that develop lupus symptoms and proteinuria spontaneously and show gender bias in disease incidence, similar to human SLE patients ^17,31-33^, have been widely used for understanding the disease etiology. We have shown that significant amounts of circulating autoantibodies against nucleohistone and dsDNA are detectable in these mice by 16 weeks of age and severe nephritis indicated by high proteinuria is observed after 20 weeks of age ^13,17^. While the timing of detectable levels of serum autoantibodies as well as proteinuria are highly heterogeneous in these mice, about 80% of the female and 20% of the male SNF1 mice develop severe nephritis within 32 weeks. We have reported that gut mucosa of female SNF1 mice, as compared to their male counterparts, harbor higher frequencies of activated B cells including plasma cells, as well as express high levels of pro-inflammatory cytokines as early as at juvenile age (4 weeks) and have amplified levels of these factors at adult ages ^17^.

In the current study, we assessed if the degree of IgA production in the gut relates to the pro-inflammatory immune phenotype of the gut mucosa. Examination of the B cells revealed that SNF1 mice have relatively higher occurrence of total and IgA+ B cells in the intestinal compartment compared to lupus resistant B6 mice (Fig. 1A). Further, IgA+ B cell frequencies, in the intestinal mucosa particularly, were relatively higher in SNF1 females compared to their male counterparts. To assess if the overall amounts of IgA produced in gut mucosa is different in lupus -prone and -resistant mice, total immune cell preparations from the intestinal tissues were tested for spontaneous IgA release. As observed in Fig. 1B, total IgA secreted by the small and large intestinal immune cells was significantly higher in SNF1 mice compared to that of B6 mice. As compared to that of B6 females, cultures of gut associated immune cells of SNF1 mice showed relatively higher age-dependent elevated production of IgA. Overall, these observations indicate that the higher degree of IgA production from the gut mucosa of lupus-prone SNF1 mice, females particularly, is reflective of the pro-inflammatory immune phenotype of their gut mucosa, which was described in our recent reports ^16,17^.

**Figure 1:**
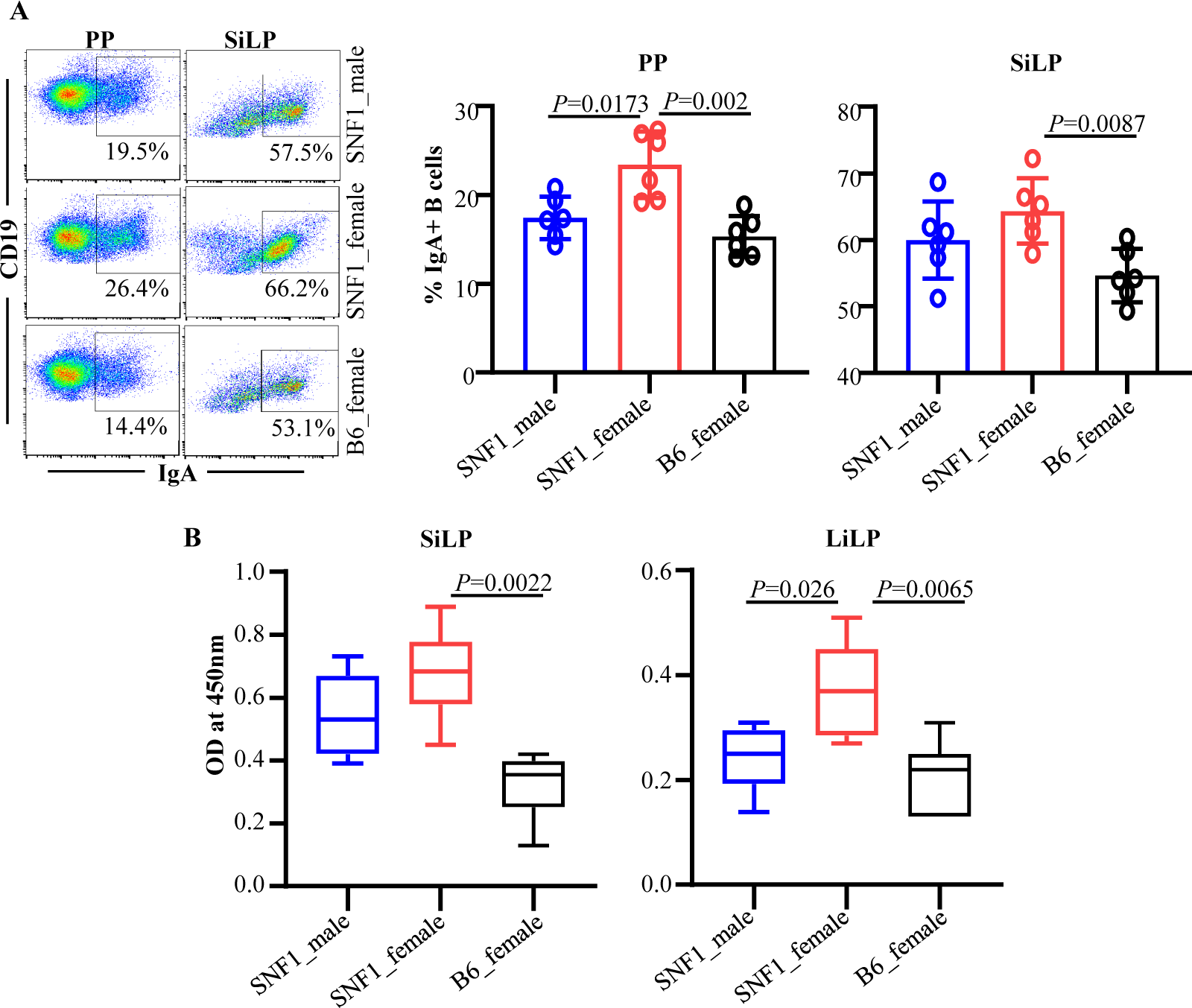
IgA+ B cell frequency and IgA secretion in the intestinal mucosa. Single cell suspensions from Peyer’s patch (PP) and small intestinal lamina propria (SiLP) were prepared from 16 week old SNF1 male, SNF1 female, and B6 female mice. **A)** These cells were subjected to FACS after staining with anti-mouse CD19 and IgA antibodies. Gating strategy for different tissues is shown in supplemental Fig. 1. Representative FACS plots (left) and mean±SD values (6 mice/group) of IgA+ B cell frequencies (right) are shown. **B)** Single cell suspensions (5×10^6^ cells/ml) were cultured for 24 hours and supernatants were tested for spontaneously secreted total IgA levels by ELISA. Mean±SD of OD values (6 mice/group) are shown. *P*-value by Mann-Whitney test.

### 3.2 IgA produced by the gut mucosa shows nAg reactivity

Since cultures of gut mucosa associated immune cells from lupus-prone SNF1 mice showed higher IgA levels compared to that of B6 mice, we examined if IgA produced in the gut mucosa of juvenile and adult age pre-nephritic mice reacts with nAg, dsDNA and nucleohistone, major lupus associated autoantigens. SiLP and LiLP cells were cultured and the supernatants were examined for nAg reactive IgA levels. As observed in Fig. 2, spent media from gut associated immune cell cultures of both juvenile and adult SNF1 mice had higher dsDNA and nucleohistone reactive IgA compared to that of B6 mice. Importantly, dsDNA and nucleohistone reactivity of IgA in the culture of gut associated immune cells from female SNF1 mice were significantly higher compared to that of their male counterparts at both juvenile and adult ages. Overall, these observations show that IgA produced in the gut mucosa of lupus-prone mice, as early as juvenile age, can recognize nAg and the degree of nAg reactivity correlates with the known gender bias of lupus incidence in this mouse model.

**Figure 2:**
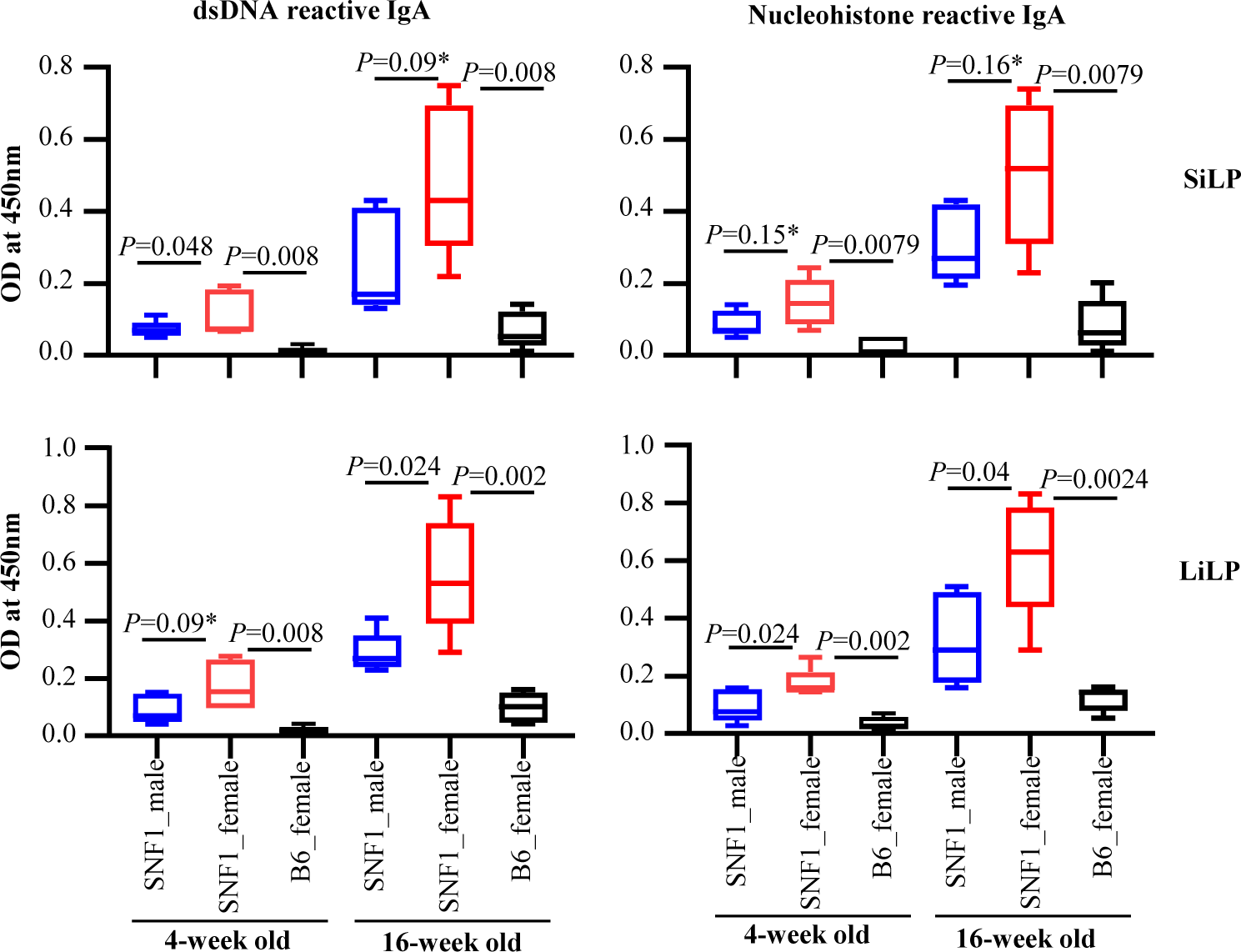
Nuclear antigen (nAg) reactivity of IgA produced by the gut mucosa cells. Immune cell rich fractions were prepared from single cell suspensions of small and large intestinal lamina propria (SiLP and LiLP) of 4 and 16 week old SNF1 males, SNF1 females and B6 females by Percoll gradient method. These cells were cultured (2×10^6^ cells/ml) for 48h in the presence of bacterial LPS and anti-CD40 antibody. Supernatants were tested for dsDNA and nucleohistone reactive IgA levels by ELISA. Mean±SD of OD values (5 mice/group) are shown. *P*-value by Mann-Whitney test.

### 3.3 SNF1 mice show higher abundance of IgA in the feces

Since gut associated immune cell cultures showed higher levels of IgA antibody, longitudinal fecal samples from lupus-prone male and female SNF1 mice and age matched female B6 mice were examined for IgA levels. Quantification of IgA in fecal samples collected from 16 mice/group at different ages revealed that fecal levels of IgA are significantly higher in male and female lupus-prone SNF1 mice at all ages, including juvenile, compared to that of female B6 mice (Fig. 3A). Further, differences in the fecal IgA levels in lupus-prone SNF1 mice and lupus-resistant B6 mice were more pronounced in older adults compared to juveniles. Importantly, compared to male counterparts, female SNF1 mice showed relatively higher amounts of fecal IgA as early as at juvenile age. Of note, fecal samples collected from a cohort of female Balb/c mice (lupus-resistant) at the age of 8 weeks showed significantly lower IgA levels compared to that of age-matched SNF females (not shown).

**Figure 3:**
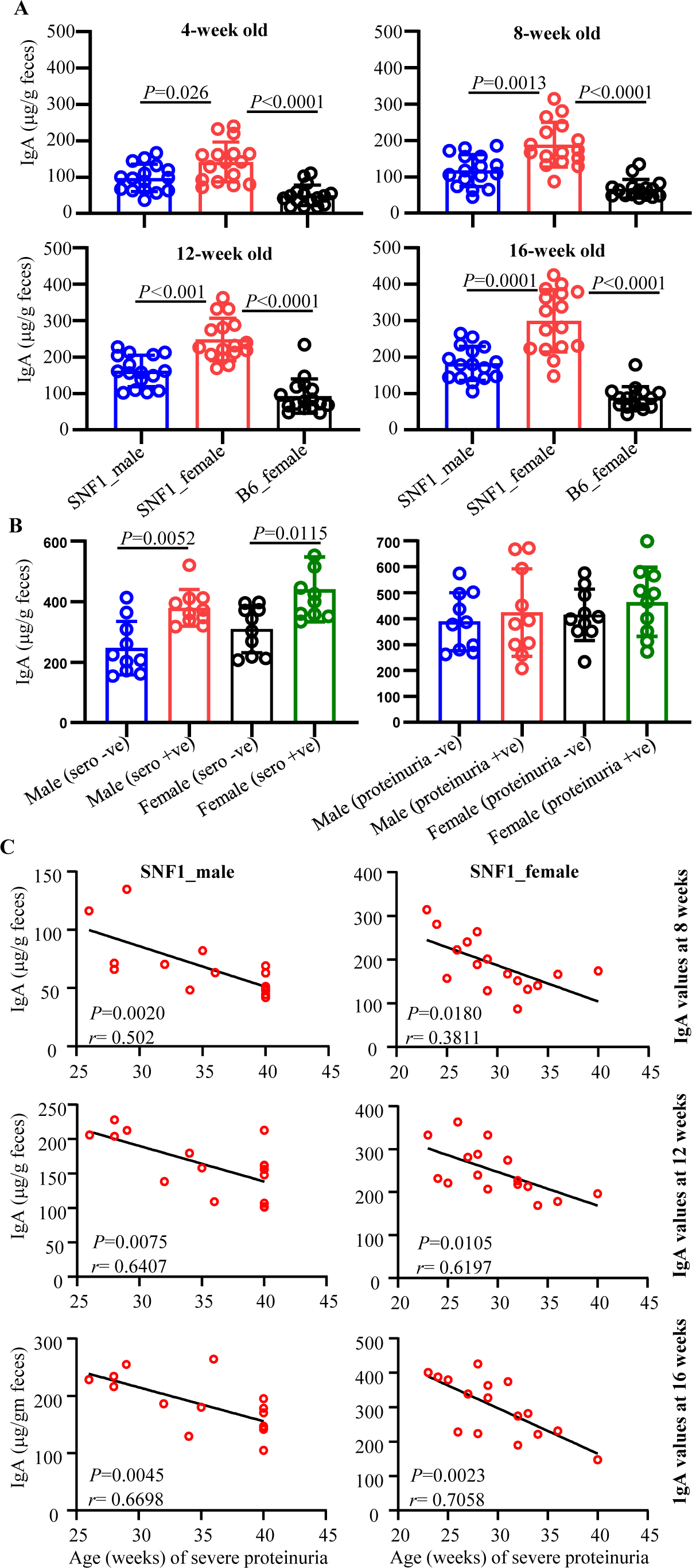
Abundance of IgA in the feces of lupus-prone and -resistant mice. Fecal samples were collected from individual SNF1 males, SNF1 females and B6 females at 4, 8, 12 and 16 weeks of age. A) Fecal pellets were subjected to ELISA to determine the IgA concentrations as detailed in Materials and method. Mean±SD of OD values (16 mice/group) are shown for each time point. B) Fecal samples from an independent cohort of age-matched seropositive and seronegative mice (between 16 and 24 weeks of age) and seropositive and proteinuria positive and seropositive and proteinuria negative (between 24 and 35 weeks of age) were also subjected to ELISA for IgA levels. Mean±SD of IgA concentration per gram feces (10 mice/group) are shown. **C)** Mice described for panel A were monitored for proteinuria levels for up to 40 weeks of age and the timing of severe proteinuria (protein: >5 mg/ml) onset was correlated with the 8-, 12- and 16-week fecal IgA levels of individual mice by Pearson’s correlation approach.

It is not known if systemic autoantibody profile and disease activity correlates with fecal IgA levels. Hence, using fecal samples from independent cohorts of age matched mice, we examined if the fecal IgA levels in seronegative and seropositive as well as in seropositive and proteinuria positive SNF1 males and females are different. Since aforementioned 16 mice/group, that were followed longitudinally, were not sufficient to obtain age matched seropositive and seronegative and age matched seropositive and proteinuria positive mice at the same time, cross-sectional samples (10 mice/group) were obtained from a much larger scale experiment. Fig. 3B shows that both male and female seropositive SNF1 mice showed significantly higher levels of fecal IgA compared to their seronegative counterparts. Similarly, although not significant compared to that of seropositive mice, higher fecal IgA levels were observed in seropositive and proteinuria positive SNF1 male and females compared to seronegative counterparts. Importantly, many of the seronegative male and female mice showed higher levels of fecal IgA suggesting that higher IgA production in the gut precedes systemic autoantibody production.

Since Fig. 3B suggested higher fecal IgA levels before sero-positive stage, we correlated fecal IgA levels of samples collected from individual pre-nephritic mice at 12 week and 16 week of age (longitudinal samples described for Fig. 3A) with the timing of proteinuria onset in them. As observed in Fig. 3C, significant correlation between their younger age fecal IgA levels and eventual clinical stage of disease is seen in both male and female mice. Overall, these observations show a positive correlation between fecal IgA levels of pre-seropositive younger ages and the eventual disease onset in SNF1 mice, and suggest that fecal IgA levels could serve as a marker for systemic autoimmune progression of pre-clinical stages.

### 3.4 Anti-dsDNA and nucleohistone reactive fecal IgA antibodies are detected in SNF1 mice

Since fecal IgA levels were higher in lupus-prone SNF1 mice compared to B6 mice and in SNF1 females compared to male counterparts, we determined if the fecal IgA antibodies from SNF1 mice recognize nuclear antigens such as dsDNA and nucleohistones, and this feature correlates with systemic autoimmunity and disease incidence. Both longitudinal and cross-sectional samples described for Fig. 3 were used for these analyses. Analysis of longitudinal samples from 16 mice/group showed that, compared to age-matched female B6 mice, both male and female SNF1 mice showed significantly higher levels of dsDNA and nucleohistone reactive IgA antibodies in their fecal samples at all ages (Fig. 4A). dsDNA and nucleohistone reactivity of fecal IgA were detectable primarily in female SNF1 mice at juvenile (4 weeks of) ages. Further, although fecal IgA antibodies of majority of the male and female SNF1 mice showed dsDNA and nucleohistone reactivity at older ages, greater number of female SNF1 mice had significantly higher levels of nAg-reactive fecal IgA compared to their male counterparts at all tested adult ages. Notably, fecal samples collected from 8 week old female Balb/c mice, similar to age matched B6 mice, showed signficantly lower nAg reactivity compared to that from SNF1 females (not shown).

**Figure 4:**
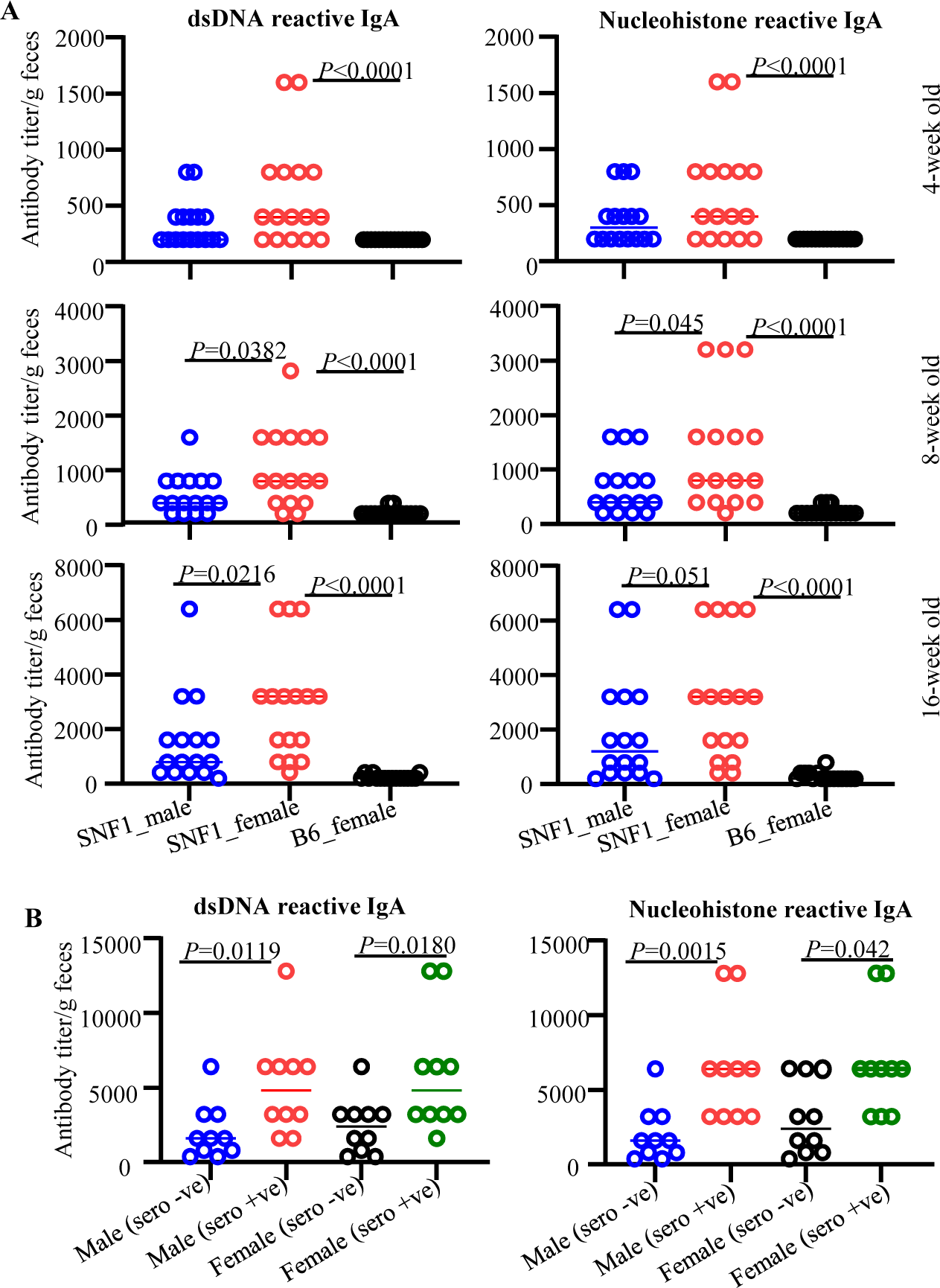
Anti-dsDNA and nucleohistone reactive fecal IgA antibodies in SNF1 mice. Fecal samples were collected from individual SNF1 males, SNF1 females and B6 females at 4, 8 and 16 weeks of age. A) Fecal pellets were subjected to ELISA to determine anti-dsDNA and nucleohistone reactivity as detailed in Materials and method. Mean±SD of ELISA titer per gram feces (16 mice/group) are shown for each time point. B) Fecal samples from an independent cohort of age-matched seropositive and seronegative mice (between 16 and 24 weeks of age) were also subjected to ELISA for anti-dsDNA and nucleohistone IgA levels. Mean±SD of ELISA titer per gram feces (10 mice/group) are shown.

As described for total IgA analysis of Fig. 3B, cross-sectional samples of age-matched sero-negative and seropositive male and female mice (10/group) were used for comparing anti-nAg IgA levels in age-matched seronegative and seropositive SNF1 males and females. As shown in Fig. 4B, nAg reactivity titers of fecal IgA were higher in seropositive male and female SNF1 mice compared to their seronegative counterparts. This analysis shows that nAg reactive IgA antibody can be found in fecal samples long before the lupus associated autoantibodies are detectable in the systemic compartment.

Since Fig. 4B suggested higher nAg reactive fecal IgA levels before sero-positive stage in both male and female SNF1 mice, we correlated fecal nAg reactive IgA levels of samples collected from individual pre-nephritic mice at 8- and 16-weeks of age with the timing of proteinuria onset in them. As observed in Fig. 5, a strong correlation between early-age (8 weeks; Fig. 5A and 16 weeks; Fig. 5B) nAg reactivity of both male and female SNF1 mice with the timing of eventual onset of proteinuria was detected. These observations suggest that nAg reactivity of fecal IgA at pre-seropositive stages could be predictive of the eventual disease onset in SNF1 mice.

**Figure 5:**
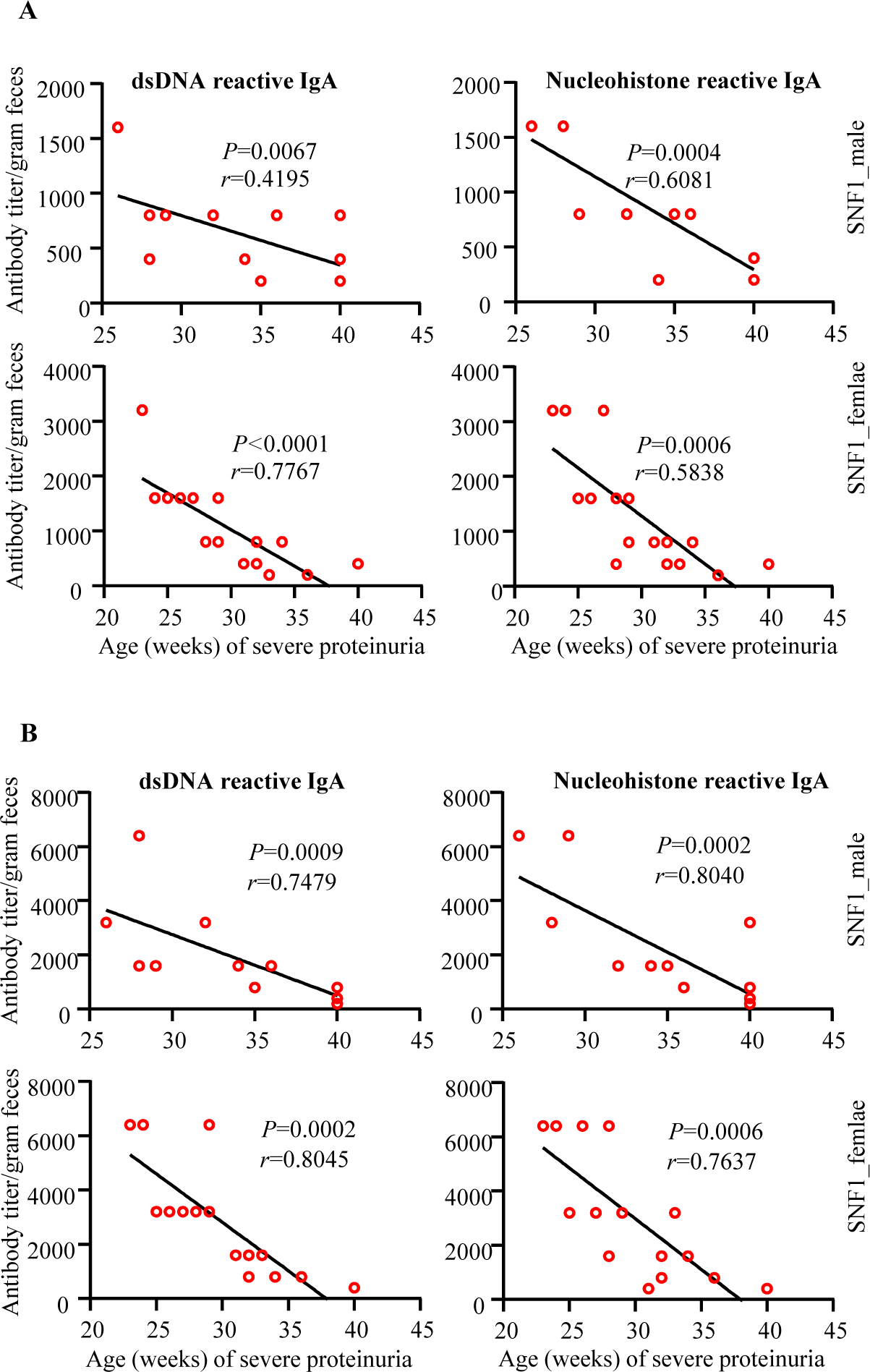
Pre-clinical stage anti-dsDNA and nucleohistone reactive fecal IgA levels correlate with eventual proteinuria onset in SNF1 mice. Mice described in Fig.4 A were monitored for proteinuria levels for up to 40 weeks of age and the timing of severe proteinuria (protein: >5 mg/ml) onset was correlated with the 8- and 16-week fecal anti-dsDNA and nucleohistone IgA levels of individual mice by Pearson’s correlation approach.

### 3.5 Depletion of gut microbiota results in suppression of fecal IgA abundance and nAg reactivity

Microbes are the primary trigger of IgA production in the gut mucosa ^34,35^. Our recent study showed that depletion of gut microbiota suppresses the intestinal pro-inflammatory immune phenotype at juvenile and adult ages, systemic autoimmune progression and proteinuria onset in female SNF1 mice. Therefore, we examined if IgA production in the gut mucosa, and abundance and nAg reactivity of fecal IgA are influenced by gut microbiota, by treating female SNF1 mice with broad spectrum antibiotic cocktail (Abx). As reported in our recent study^16^, Abx treatment effectively depleted majority of the gut microbiota and caused a profound delay in proteinuria onset (supplemental Fig. 2). Importantly, as observed in Fig. 6A, IgA+ B cell frequencies were significantly lower in microbiota depleted mice compared to untreated mice. Further, intestinal immune cell cultures from microbiota depleted mice showed significantly lower amounts of spontaneously released IgA antibodies compared to that of untreated control mice (Fig. 6B). Cultures of intestinal immune cells from microbiota depleted mice showed significantly lower nAg reactive IgA levels compared to that of control mice (Fig. 6C). Correspondingly, Fig. 6D shows that microbiota depletion caused significant reduction in IgA abundance in fecal samples. Moreover, dsDNA and nucleohistone reactive fecal IgA levels were significantly lower in female SNF1 mice with depleted microbiota compared to their counterparts with intact gut microbiota (Fig. 6E). Of note, we have shown that serum samples from microbiota depleted SNF1 mice have significantly lower anti-dsDNA and nucleohistone IgG antibodies compared to mice with intact microbiota and the disease onset is delayed in them upon microbiota depletion^16^. Overall, these observations indicate that fecal IgA abundance and their nAg reactivity in lupus-prone SNF1 mice are, at least in part, microbiota dependent and validate that these features of fecal IgA are reflective of the degree of systemic autoimmune activity.

**Fig. 6:**
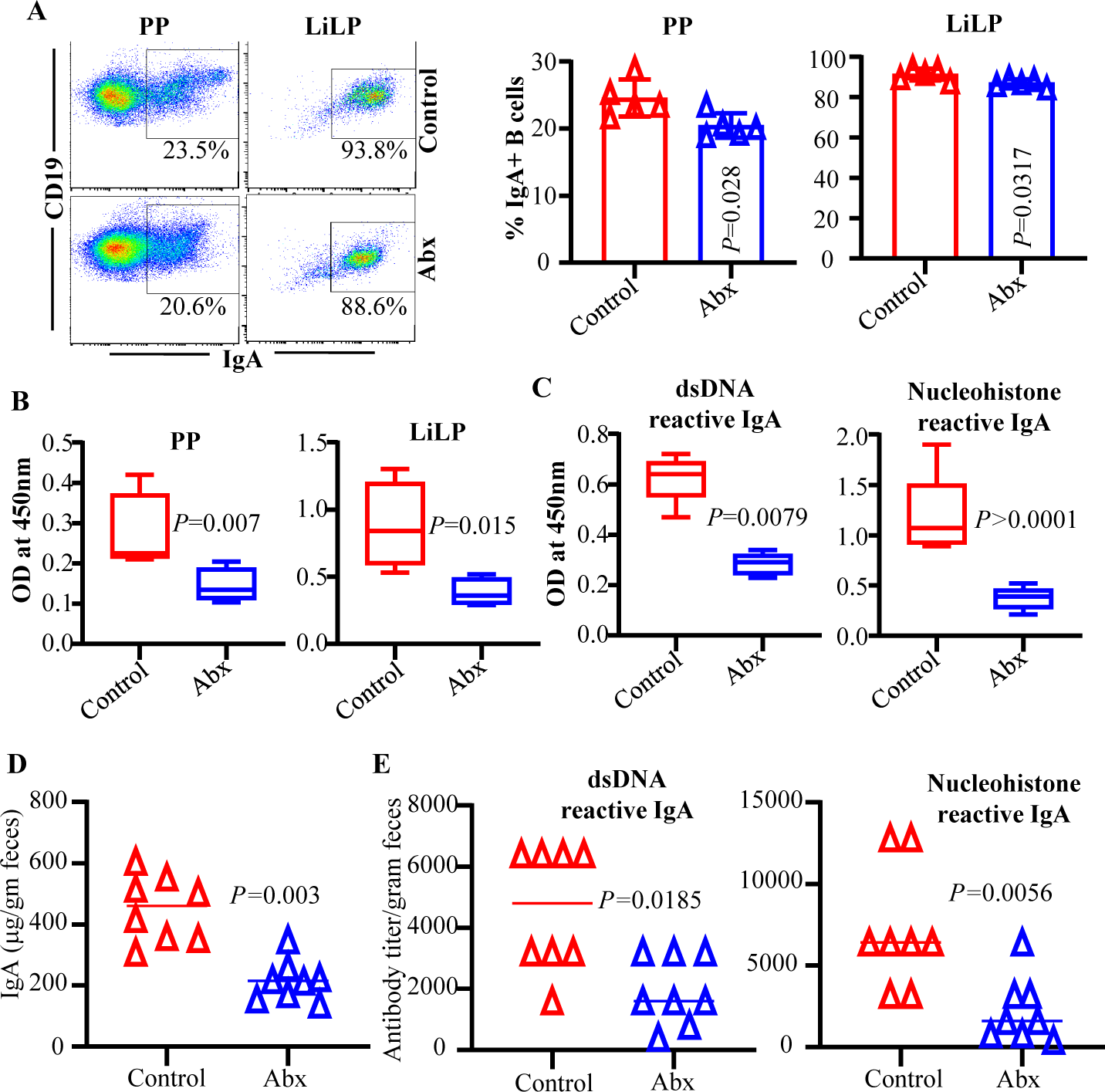
Fecal IgA abundance and nAg reactivity in microbiota depleted mice. SNF1 females were left untreated or given broad spectrum antibiotic cocktail to deplete the gut microbiota as depicted in supplemental Fig. 2A. Microbiota depletion efficacy and proteinuria monitoring data are shown in supplemental Fig. 2B and 2C. A) PP and LiLP cells from these mice at 16 weeks of age were subjected to FACS after staining using anti-mouse CD19 and IgA antibodies as described for Fig. 1. Representative FACS plots (left) and mean±SD values (5 mice/group) of IgA+ B cell frequencies (right) are shown. B) PP cells and immune cell rich fractions of LiLP were cultured (2×10^6^ cells/ml) for 48h in the presence of bacterial LPS and anti-CD40 antibody. Supernatants were tested for total IgA levels by ELISA. C) Supernatants of LiLP cultures were tested for dsDNA and nucleohistone reactive IgA levels by ELISA. Mean±SD of OD values (5 mice/group) are shown in panels B & C. D & E) Fecal pellets from another cohort of control and microbiota depleted mice (20 weeks of age) were subjected to ELISA to determine the total IgA, and dsDNA and nucleohistone reactive IgA levels. Mean±SD of OD values (8 mice/group) are shown. *P*-value by Mann-Whitney test.

## 4. Discussion

Immunoglobulin A not only neutralizes the pathogens at mucosal surfaces, but also regulates the composition and function of gut microbiota by stabilizing the intestinal colonization by symbiotic microorganisms ^20-22^. Recent reports have shown that the degree of IgA production in the gut mucosa under normal and clinical conditions including SLE can be different ^9,36,37^. A recent report has shown that fecal IgA and IgG levels are higher in SLE patients compared to healthy controls ^37^. Nevertheless, if IgA produced in the gut mucosa contributes to autoimmune process or higher antibody production in the gut mucosa in SLE patients precedes systemic autoimmunity and clinical onset of the disease or is the consequence of autoimmune process is unknown. Further, an association between IgA production in the gut mucosa and gender bias associated with lupus has not been studied. Our recent reports ^16,17^ showed, for the first time, that the immune phenotype of gut mucosa is significantly different in lupus-prone male and female SNF1 mice. Pro-inflammatory immune phenotype of gut mucosa including higher frequencies of plasma cells appear in female SNF1 mice at juvenile age, much before the production of sex hormones and detectable levels of circulating autoantibodies^17^. These observations suggest that the production of IgA in gut mucosa and its levels in feces, may be significantly different in males and females under lupus susceptibility prior to disease onset. In the current study, we detected higher amounts of IgA antibodies in the fecal samples of lupus-prone SNF1 mice compared to lupus-resistant B6 mice and found that the levels of these antibodies correlate positively with rapid disease progression and higher disease incidence of female SNF1 mice. Most importantly, we also found, for the first time, that nuclear antigen-recognizing fecal IgA antibodies can be detected in lupus-prone mice as early as juvenile age and these fecal antibody levels at younger ages correlate with eventual circulating autoantibody levels and disease progression at adult ages, and the gender.

SNF1 mice that develop lupus symptoms and proteinuria spontaneously and show strong gender bias in disease incidence, similar to human SLE patients ^17,31-33^, have been used for understanding the disease etiology. Previously, we have shown that significant amounts of circulating autoantibodies against nucleohistone and dsDNA are detectable in these mice by 16 weeks of age and severe nephritis indicated by high proteinuria after 20 weeks of age also develops ^13,17^. While the timing of detectable levels of serum autoantibodies as well as proteinuria are heterogeneous, about 80% of the female and 20% of the male SNF1 mice develop severe nephritis within 32 weeks. Hence, our observation that IgA levels, although highly heterogeneous, are higher in significant number of lupus-prone females compared to males even at juvenile age, much before the clinical onset of disease, is noteworthy. Importantly, fecal levels of IgA at younger ages showed excellent correlation with older adult age proteinuria onset indicating that fecal IgA levels may serve as biomarker for early detection of clinical onset of SLE. These observations, along with our recent reports showing pro-inflammatory immune phenotype of gut mucosa including higher amounts of large number of pro-inflammatory cytokines and plasma cells that appear in female SNF1 mice at juvenile age ^16,17^, indicates that pro-inflammatory cytokines and IgA produced in the gut mucosa may be involved in the initiation and perpetuation of systemic autoimmunity in lupus.

It is well established that gut microbiota including symbionts in general, and specific microbial communities including pathobionts in particular, can influence the magnitude of antibody production in the gut mucosa and the IgA abundance in feces ^34,35,38-40^. Interestingly, however, our recent report showed that gut microbiota composition is significantly different in lupus-prone males and females only at adult age, but not at juvenile age ^16^, suggesting that the production of higher pro-inflammatory cytokines and IgA in the gut mucosa at younger ages is not due to differences in the gut microbiota. It is possible that differences in the types and degrees of host-microbe molecular interactions at gut mucosa may be responsible for differences in the immune phenotypes of lupus-prone males and females. Irrespective of the mechanisms involved, pro-inflammatory features including hyper-activation of B cells of the gut mucosa in the presence of gut microbial components could aid in the production of anti-microbial antibodies that cross react with host antigens. In this regard, microbial antigens are continuously sampled by immune cells of the gut mucosa and it can result in local and systemic responses ^41-44^. Further, host-microbial interactions could aid in autoimmune initiation and progression in at-risk subjects, via multiple mechanisms including bystander activation and molecular mimicry ^45-47^. In fact, contribution of molecular mimicry, primarily by antigens of pathogenic bacteria, in the initiation and/or perpetuation of T and B cell responses to self-antigens have been widely investigated ^48,49^. A recent report has shown that pathogenic autoreactive T and B cells can cross react with mimotopes expressed by a human gut commensal and contribute to autoimmunity in an anti-phospholipid syndrome (APS) model ^50^.

Autoantibody response against antigens such as nucleic acids and nuclear proteins including histones, is the key feature of lupus autoimmunity in humans and rodent models. Importantly, nucleic acids and histone-like and other nucleic acid binding proteins can be structurally homologous to their mammalian host counterparts ^51-55^, suggesting that cross-reactive antibodies against microbial and host molecules are generated in the gut mucosa. This also prompts the notion that antibody production in the gut mucosa against these microbial components could eventually spread to the systemic compartment in lupus-prone background as autoimmunity against host nuclear antigens. It is possible that pro-inflammatory gut mucosa in a lupus-susceptible background, as we observed in lupus-prone mice^16,17,56^, could facilitate this process. In fact, our observation that fecal IgA in lupus-prone female SNF mice are dsDNA and nucleohistone reactive as early as juvenile age, much earlier than the appearance of circulating autoantibodies, supports this notion. Importantly, age-dependent progressive increase in the dsDNA and nucleohistone reactive fecal IgA and strong correlation between the levels of these antibodies at younger ages and the eventual systemic autoimmune progression and proteinuria was observed in this study. This suggests a vicious cycle of pro-inflammatory responses in the gut mucosa contributing to the generation of a large number of host nuclear antigen and microbial antigen cross-reactive B cells in the gut mucosa and their eventual spread to the systemic compartment leading to the pathogenic events of systemic autoimmunity in lupus.

IgA-antibody production in the gut mucosa is largely influenced by microbiota and the germ free (GF mice) as well as microbiota-depleted SPF mice show significantly lower amounts of fecal IgA ^38,57-59^. Gut microbial communities including symbionts and pathobionts could differently impact the degree of IgA production in the gut mucosa ^35,60-62^. Therefore, we postulated that if anti-nuclear antigen reactive IgA response in the gut mucosa of lupus-prone mice is originated against microbial antigens, then depletion of gut microbiota could suppress nAg reactive IgA levels. Of note, recently we showed that the depletion of gut microbiota using broad spectrum antibiotics suppressed autoimmunity and gender bias, by causing slower disease progression in SNF1 females ^16^. Further, another group has reported that selective depletion of gut microbiota using vancomycin suppresses systemic autoimmunity and serum IgA abundance in lupus-prone MRL/lpr mice ^63^. Here we show that depletion of gut microbiota not only caused the suppression of fecal total IgA levels, but also affected the degree of dsDNA and nucleohistone reactivity.

Overall, our study demonstrates, for the first time, that fecal IgA profiles are different in lupus-prone SNF1 mice and lupus-resistant B6 mice as early as juvenile age and this feature is reflective of late adult age systemic autoimmunity and nephritis. Most importantly, we also found that fecal IgA in lupus-prone mice show nAg reactivity at younger ages, long before the detection of systemic autoantibodies and lupus nephritis. Furthermore, fecal IgA abundance and nAg reactivity correlate with the onset of lupus-like disease associated gender bias in this mouse model. Of note, (although not shown) fecal pellets from a single age group of Balb/cJ females from Jackson lab also showed lower total and nAg reactive fecal IgA levels compared to that of SNF counterparts. However, additional systematic longitudinal comparative studies are needed using different age groups of Balb/cJ and other lupus-resistant strains of mice that are housed and bred in our colony to realize the strain to strain difference in fecal IgA features including nAg reactivity. In this regard, a recent reports showed that, as compared to B6 mice, Balb/c mice of three different vendors (Taconic, Charles River and Harlan), mice that harbor segmented filamentous bacteria particularly, produce higher amounts of polyreactive IgA in the gut mucosa and fecal samples^64^. Nevertheless, our systematic comparison between lupus-prone SNF1 mice and lupus-resistant B6 mice show a higher abundance and nAg reactivity of fecal IgA in lupus-prone SNF1 mice. Most importantly, we also found that fecal IgA in lupus-prone mice show nAg reactivity at younger ages, long before the detection of systemic autoantibodies and lupus nephritis. Furthermore, fecal IgA abundance and nAg reactivity correlate with the onset of lupus-like disease associated gender bias in this mouse model. These novel observations suggest that fecal IgA levels and nuclear antigen reactivity could serve as an early biomarker to detect the systemic autoimmune activities in at-risk subjects, before the seropositive stage and disease onset. Since the abundance and nAg reactivity of fecal IgA in lupus-prone subjects have not been studied, systematic longitudinal follow-up studies are needed to determine the clinical translation value of IgA features as biomarkers to predict the clinical disease.

## Abbreviations

SLE: systemic lupus erythematosus
SNF1 mice: (SWRxNZB)-F1 mice
nAg: nuclear antigen
PP: Peyers’ patch
LP: Lamina Propria
SI: small intestine
GF: germ-free
SPF: specific pathogen free
TLR: toll-like receptors
Abx: antibiotic cocktail
MUSC: Medical University of South Carolina
ELISA: Enzyme-linked immunosorbent assay
dsDNA: double stranded DNA

## *Acknowledgments

This work was supported by internal funds from MUSC, National Institutes of Health (NIH) grants R21AI136339 and R01AI138511. W.S. performed experiments and reviewed the manuscript, R.G. performed experiments and reviewed the paper, B.M.J. performed sample collection and reviewed the paper, and C.V. designed the study, performed the experiments, and wrote the paper. W.S. and R.G. contributed equally to this work. Dr. Vasu is the guarantor of this work and, as such, had full access to all the data in the study and takes responsibility for the integrity and accuracy of the data and analysis. The authors are also thankful to flow cytometry core of MUSC for the instrumentation support.

## Conflict of Interest statement

Authors do not have any conflict(s) of interest to disclose.

## Notes

### Competing Interest Statement

The authors have declared no competing interest.

