## Supplemental data for "Abundance and nuclear antigen reactivity of intestinal and fecal Immunoglobulin A in lupus-prone mice at younger ages correlate with the onset of eventual systemic autoimmunity"

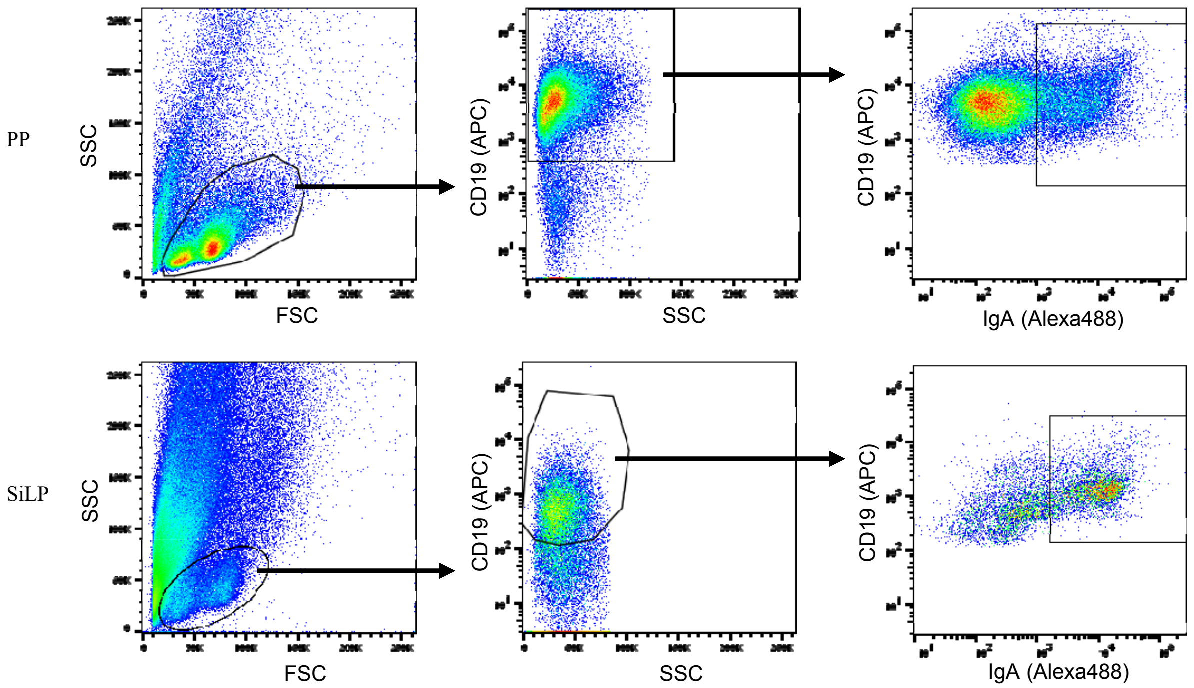


*Supplemental Fig 1:* Gating strategy employed for figures 1A. Because different cell numbers were used for staining of PP and intestine samples and the staining intensities were different due to difference in the cellular compositions, slightly different gating strategies were employed for these two sample types. Similar strategies were employed for Fig. 6A.

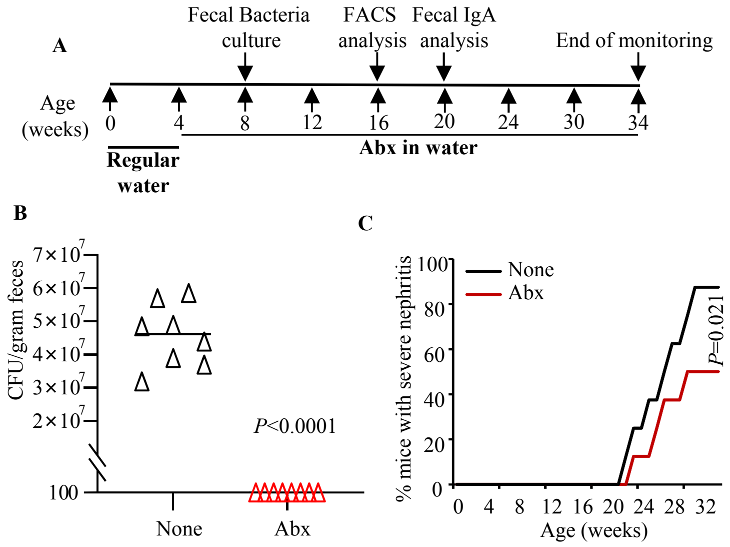


*Supplemental Fig 2:* Depletion of gut microbiota using antibiotics. Female SNF1 mice were given a broad-spectrum antibiotic cocktail [Abx: ampicillin (1 g/l), vancomycin (0.5 g/l), neomycin (1 g/l), and metronidazole (1 g/l)] -containing drinking water starting at 4 weeks of age. **A)** Schematic of treatment and time-points of various assays. **B)** Fecal pellets collected from individual treated and untreated controls at 8 weeks of age were suspended and diluted in sterile PBS, plated onto BHI medium agar plates under anaerobic and aerobic conditions for up to 72 h, the total number of colonies were counted, and colony forming units (CFU)/gram initial fecal material were calculated. n= 8 mice/group. *P*-values by two-sided Mann Whitney test. Cohorts of mice from a parallel experiment were euthanized at 16 weeks of age for immunological assays ( shown in Fig. 6A-5C). **C)** Cohorts of 8 mice/group were monitored for proteinuria for up to 34 weeks of age. *P*-values by log-rank test. Fecal pellets collected at 20 weeks of age from these mice were tested for IgA features (shown in Fig. 6D&E).
